# Machine learning prediction of eukaryotic hosts for giant viruses

**DOI:** 10.64898/2026.09.08.750034

**Authors:** Hsin-Ying Chang, Frederik Schulz, Chuan Ku

## Abstract

Giant viruses (GVs; *Nucleocytoviricota*) infect diverse eukaryotes and are ecologically important across ecosystems. Although cultivation-independent sequencing has recovered tens of thousands of GV genomes from environmental samples, eukaryotic hosts are only known for a few isolates, leaving the host contexts of most GVs elusive. We developed GVHoP (<u>G</u>iant <u>V</u>irus-<u>Ho</u>st <u>P</u>redictor) that predicts eukaryotic hosts from GV genomes by integrating gene content and GV-eukaryotic sequence similarities. Trained on isolates with experimentally identified hosts, GVHoP achieved 97% accuracy in cross-validation and predicts hosts at three hierarchical levels of eukaryote classification. Functional analyses link top predictive features to various processes involved in virus-host interactions, including viral entry, replication, morphogenesis, and cellular metabolic reprogramming. Applying GVHoP to 7, 897 GV metagenome-assembled genomes assigned hosts to 5, 280 viruses and uncovered potential novel hosts across major GV orders that cannot be inferred from core-gene phylogenetic information. The predicted host composition clearly separates aquatic and terrestrial environments and distinct aquatic ecosystems. Together, GVHoP links viruses known only from nucleic acid sequences to their putative hosts across the eukaryotic tree of life, improves our understanding of GV-host interactions and their potential impacts on ecosystems, and paves the way for further ecological and functional studies of environmental GVs.

## INTRODUCTION

Giant viruses (GVs) are nucleocytoplasmatic large DNA viruses in the viral phylum *Nucleocytoviricota*. These viruses are characterized by their large virion sizes, which range from 100 to 2, 500 nm (1, 2). Some well-known representatives, such as *Mimivirus*, *Pandoravirus*, and *Pithovirus*, are comparable to the size of a typical bacterial cell and possess complex genomes with up to 2.5 Mbp (3). They are widely distributed across global ecosystems, including marine and freshwater environments, terrestrial soils, and even extreme habitats (4–7). Isolated viruses from each clade of *Nucleocytoviricota* are known to have different, sometimes overlapping host ranges. *Algavirales* infect algae, *Chitovirales* infect animals, *Mycodnavirales* infect fungi, whereas *Imitervirales*, *Pandoravirales*, *Asfuvirales*, and *Pimascovirales* infect diverse microeukaryotes especially amoebozoans, or animal hosts (8, 9). Beyond their ecological roles in driving global nutrient and carbon cycling by regulating algal and protist populations (e.g., bloom-collapsing coccolithoviruses) (10), several GVs are critical pathogens for humans (Variola, Mpox), livestock (African swine fever virus), and aquaculture (iridoviruses) (11–14). Previous studies based on gene transfer analyses suggest that the host diversity of GVs remains underexplored (5, 15). Dated phylogenies further indicate that GV diversification involved extensive host shifts, including at least two transitions from amoebozoan to animal hosts that eventually gave rise to iridoviruses and African swine fever viruses (16). Collectively, these studies highlight the ecological and societal importance of GVs infecting diverse eukaryotes, yet most known virus-host associations are derived from isolated GVs and cultured hosts. Recent advances in metagenomic approaches have enabled high-resolution recovery of GV metagenome-assembled genomes (GVMAGs) from diverse environments such as oceans, soils, and permafrost, substantially expanding our understanding of GV diversity across ecosystems (6, 17–19). Despite these breakthroughs, a major challenge remains: nearly all of the recovered GVMAGs lack host information, limiting our ability to infer their viral biology and ecological roles.

To date, several tools have been developed to predict virus-host relationships, especially between phages and their prokaryotic hosts (20–24). Compared with most phages, GVs generally have complex mosaic gene repertoires due to extensive horizontal gene transfers (HGTs) from eukaryotes, prokaryotes, and other viruses (25–27). Similarly, their eukaryotic hosts also have much higher coding capacities than prokaryotes. Gene repertoires therefore comprise a major source of information for predicting GV-eukaryote interactions, in contrast to simpler features such as genomic *k*mer frequencies, GC content, or a limited set of markers that have been used for predicting phage-host interactions. Moreover, hosts of GVs span diverse and deeply divergent lineages of eukaryotes, especially protists (28), which presents distinct challenges from predicting phage hosts at the strain level. Although some tools have been developed for predicting the eukaryotic hosts of viruses, such as Host Taxon Predictor (HTP), VIDHOP, and EvoMIL (29–31), key limitations remain. HTP and VIDHOP use simple sequence features (e.g., *k*mer frequencies), and they can only predict a limited number of hosts, mainly targeting human or disease-related viruses. EvoMIL predicts both prokaryotic and eukaryotic hosts using protein language models and attention-based multiple instance learning, but eukaryotic prediction remains inherently difficult and is limited to animal or plant hosts (31).

In this work, we develop GVHoP, a machine learning model designed to predict the eukaryotic host taxa of GVs. Leveraging host-annotated genomic data from GV isolated genomes (GVIGs), we curated a high-confidence training dataset of 252 complete genomes with identified hosts. The GVHoP pipeline harnesses the information of GV gene content and GV-eukaryote gene similarities (hereafter referred to as GV-euk signals) to determine the most likely host of a GV genome. The model also ranks feature importance to uncover critical candidate genes linked to GV-host interactions. By applying GVHoP to GVMAGs, we further expanded our knowledge about the ecological roles of thousands of uncultivated viruses.

## MATERIAL AND METHODS

### Curated dataset of GVIGs and simulated GVMAGs for model development

Available viral genomes with known hosts were retrieved from the NCBI database and organized into a dataset covering records up to October 2025. To complement the dataset, new releases of viral genomes with known hosts were downloaded manually, including *Catovirus naegleriensis* (32) and *Naiavirus* (33). A dataset of 252 GVIGs was compiled, with accession numbers, virus taxonomy, host taxonomy, and genome sizes (Supplementary Table 1). To simulate GVMAGs, each GVIG was artificially fragmented to predefined completeness levels of 50%, 60%, 70%, 80%, and 90% with modified genome_fragmentizer.py from https://zenodo.org/records/10085666 (34).

### Phylogenetic inference and phylogenetic distance-based clustering of GVIGs

Seven GV core proteins, superfamily II helicase [SFII], large RNA polymerase subunits [RNAPL], family B DNA polymerase [PolB], TFIIB transcriptional factor [TFIIB], topoisomerase family II [TopoII]), DNA-packaging ATPase, and virus late transcription factor 3 [VLTF3] (35), were extracted using ncldv_markersearch (https://github.com/faylward/ncldv_markersearch) from CDS sequences predicted with Prodigal v2.6.3 in meta option (36). Protein sequences were aligned using Muscle5 (37), followed by removal of poorly aligned positions with trimAl v1.5 (38) using the automated setting. IQ-TREE v2.3.6 (39) was used for maximum-likelihood inference, with -m MFP setting for substitution model selection (LG+F+I+R8). Node support was assessed through 1, 000 bootstrap replicates. Closely related genomes were grouped according to phylogenetic closeness using the Phylogenetic Distance Matrix Clustering (PDMClust) approach (https://github.com/NeLLi-team/pdmclust). For downstream model development, a PDM cutoff of 0.95 was used to represent the species level within the GV phylogeny.

### Feature engineering

#### Gene content

An all-against-all sequence similarity search was performed on 98, 419 protein sequences from GVIGs using BLAST v2.6.0 (40) with an e-value cutoff of 10□□. Based on the resulting similarity scores (e-values and bit scores), proteins were clustered into orthologous gene clusters (orthogroups; OGs) using OrthoMCL v1.3 (41) with an inflation parameter of 1.1 and a maximum edge weight of 300, yielding 25, 129 OGs in total. OGs present in only a single genome (16, 836 OGs) were excluded, gene counts of the remaining OGs (8, 293 OGs) were used as feature values in the gene content matrix.

#### GV-euk signals

A eukaryotic database was derived from the NCBI non-redundant protein database and included only sequences from the supergroups Amoebozoa, Archaeplastida, Discoba, Haptista, Opisthokonta, and SAR, generating a total of 146, 333, 597 sequences. The same set of 98, 419 GVIG protein sequences was queried against the eukaryotic database using DIAMOND v0.9.24.125 (42) for fast protein alignment with an e-value cutoff of 10□□. The eukaryotic sequences without any hit (146, 172, 840 sequences) or with only one hit from a single GVIG genome (103, 507 sequences) were removed. The corresponding bit scores of the remaining 57, 250 eukaryotic proteins were used as feature values in the GV-euk signal matrix.

#### kmer frequency

To profile local sequence composition across viral genomes, *k*mers (contiguous substrings of length *k*) were extracted using a custom Python script adapted from VirHostMatcher (22) (<u>repository: VirHostMatcher-Net</u>). The frequency of each *k*mer substring was calculated to construct a feature matrix.

### Architecture of classification models

#### Base-classifiers

An eXtreme Gradient Boosting (XGBoost) classifier v3.0.1 (https://github.com/dmlc/xgboost) was employed for supervised learning (43). Model hyperparameters, the number of estimators was fixed at 1, 000, while the maximum tree depth was tuned using 5-fold cross-validation to achieve optimal predictive performance. The GPU-accelerated version of XGBoost was executed on in-house computing nodes equipped with NVIDIA RTX 4090 (24GB) GPUs to ensure the computational efficiency. We applied a hierarchical classification framework in which XGBoost classifiers were trained at each level of the host hierarchy.

#### Meta-classifier

A meta-classifier based on neural network (PyTorch v2.6.0; https://pytorch.org) was built on the raw scores from each base-classifier, the top feature set from the gene content-based models, and the top feature set from the GV-euk signals-based models, which were organized into a feature matrix. The training data were standardized on a feature-wise basis using the StandardScaler (Scikit-learn v1.6.1; https://scikit-learn.org/stable/) to normalize the distribution of each input variable before it is processed by the neural network. The network architecture consisted of an input layer followed by a three-layer feedforward network: an initial 64-unit dense layer with ReLU activation, batch normalization (BN), and 50% dropout; a subsequent 32-unit dense layer with ReLU, BN, and 30% dropout; and a final 64-unit dense layer with ReLU activation. The output layer applied a *softmax* activation function to generate probability estimates for host bottom-level labels. Model optimization was performed using the Adam optimizer with a learning rate of 0.001 and categorical cross-entropy as the loss function. Early stopping based on validation loss was implemented to prevent overfitting and improve model generalization. For the models described above, the potential class imbalance within the dataset, class weights were computed and incorporated during training.

### Feature importance and functional annotation

Feature importance scores contributing to the model predictions were computed using the built-in function implemented in the model framework (.feature_importances_). The protein sequences corresponding to the top gene-content features were functionally annotated using eggNOG-mapper v2.1.3 (44) against the eggNOG database v5.0.2 (45). Functional assignments included KEGG orthologs (KOs) (46), clusters of orthologous groups (COGs) (47), and PFAM protein families (48). Foldseek was employed to search against the Protein Data Bank using predicted 3Di structural sequences generated by ProstT5 (49, 50). The top GV-euk signal features were annotated with their gene names retrieved from NCBI nr using EPost v25.9 (https://www.nlm.nih.gov/dataguide/edirect/epost.html).

### Hierarchical selective classification and coverage

#### Bottom-up approach

Following the bottom-up inference strategy (51), predictions are initiated at the most specific level of the hierarchy. The classifier produces scores for the available host classes, which are converted into class probabilities using a *softmax* function. The class with the highest probability is selected as the prediction for that level. The set of possible classes and their relationships therefore follow the predefined host hierarchy. If the predicted probability is below a predefined confidence threshold, the prediction is considered insufficiently confident. The model then moves one level upward by aggregating the probabilities of the descendant classes belonging to the same parent category. This process is repeated until the aggregated probability exceeds the confidence threshold, at which point the corresponding host category is reported as the final prediction. Hierarchical accuracy was evaluated at the level where the confidence threshold was first achieved. A prediction was considered hierarchically correct when the predicted node corresponded to the ground-truth node or to an ancestor of the ground-truth host within the predefined hierarchy. This allows predictions to retain partial hierarchical information while avoiding over-specific assignments that do not meet the required confidence level.

#### Hierarchical coverage

Hierarchical coverage is adopted to quantify the specificity of predictions within the host hierarchy (51). As predictions at more specific hierarchical levels provide more information, recent studies have introduced entropy-based measures to characterize the uncertainty associated with the descendant classes of a given hierarchical node. Let *v* denote the set of hierarchical nodes and *L*(*v*) denote the set of leaf nodes descending from node *v* ∈ *V*. Under the assumption of a uniform prior distribution over the descendant leaf nodes, each leaf node under contributes equally to the uncertainty measure. The corresponding entropy is therefore defined as:

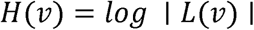

Under this formulation, entropy attains its maximum value at the root node *r*, corresponding to the least informative prediction, and its minimum value at the leaf nodes, corresponding to the most informative prediction. Based on this property, hierarchical coverage is defined as the normalized entropy reduction of node relative to the root node:

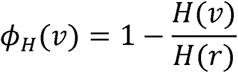

Accordingly, the root node has zero coverage because it provides no specific host information. Coverage increases as predictions move toward deeper levels of the hierarchy, reaching a maximum value of 1 at the most specific nodes.

### Principal coordinate analysis

Using SciPy (v1.15.2; https://scipy.org) pdist and squareform functions, we converted the two top feature sets, gene content and GV-eukaryote signals, into cosine distance matrices. Host composition dissimilarity across environments was quantified by converting the GVMAG-predicted host ratio matrix into a Bray-Curtis distance matrix. The downstream Principal Coordinate Analysis (PCoA) was executed using the pcoa function in Scikit-bio (v0.7.3; https://scikit.bio).

### GVMAGs database

Species-level uncultivated viral genomes (*n*=8, 508) were downloaded from the GVMAG repository (19) (https://gvmagdb.newlineages.com/downloads/file/mag-fna and https://gvmagdb.newlineages.com/downloads/file/mag-faa).

### Prediction purity of GVMAGs

The prediction purity quantifies the extent to which host assignments of GVMAGs inferred from GVIGs within the same *Nucleocytoviricota* clade are consistent with the predicted host. To compute the prediction purities, FastTree 2 (52) was used to establish a phylogeny of GVMAGs and GVIGs based on the same core proteins as aforementioned. Relative Evolutionary Distance (RED) scores were computed using the R package Castor v1.8.5 (53) represent the relative position of each node from the root to the tips, providing a measure of evolutionary divergence. The RED scores were used to cluster genomes at different levels of divergence. For an unseen GVMAG sample *Z_i_*, *N_k_*(*Z_i_*) denotes the set of the *k* nearest reference genomes (GVIGs) to *Z_i_* based on a RED-derived cluster. The predicted host of *Z_i_* is *ŷ_i_*. The prediction purity is then defined as the proportion of neighboring reference genomes belonging to the same predicted host within the *k*-nearest phylogenetic neighborhood:

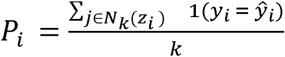

## RESULTS

### Development of GVHoP models

#### Data collection and preprocessing

To build supervised machine learning models for host prediction of GVs, we used a dataset of 252 GVIGs with host labels as training data (Supplementary Table S1, Figure 1A). A phylogeny of these GVIGs was inferred from a concatenated alignment of seven core proteins and rooted between *Megaviricetes* and *Pokkesviricetes* (Figure 2A). Consistent with the International Committee on Taxonomy of Viruses (ICTV) (2025; https://ictv.global/taxonomy) and earlier research (35, 54), these viruses are grouped into seven orders: *Algavirales*, *Asfuvirales*, *Chitovirales*, *Imitervirales*, *Pandoravirales*, *Pimascovirales*, and the newly proposed fungi-infecting *Mycodnavirales* (8). These GVIGs were also clustered using Phylogenetic Distance Matrix Clustering (PDMClust), which resulted in 36 species-level clusters (more than 2 genomes) and 85 singletons (Figure 2A, 2B).

**Figure 1.**
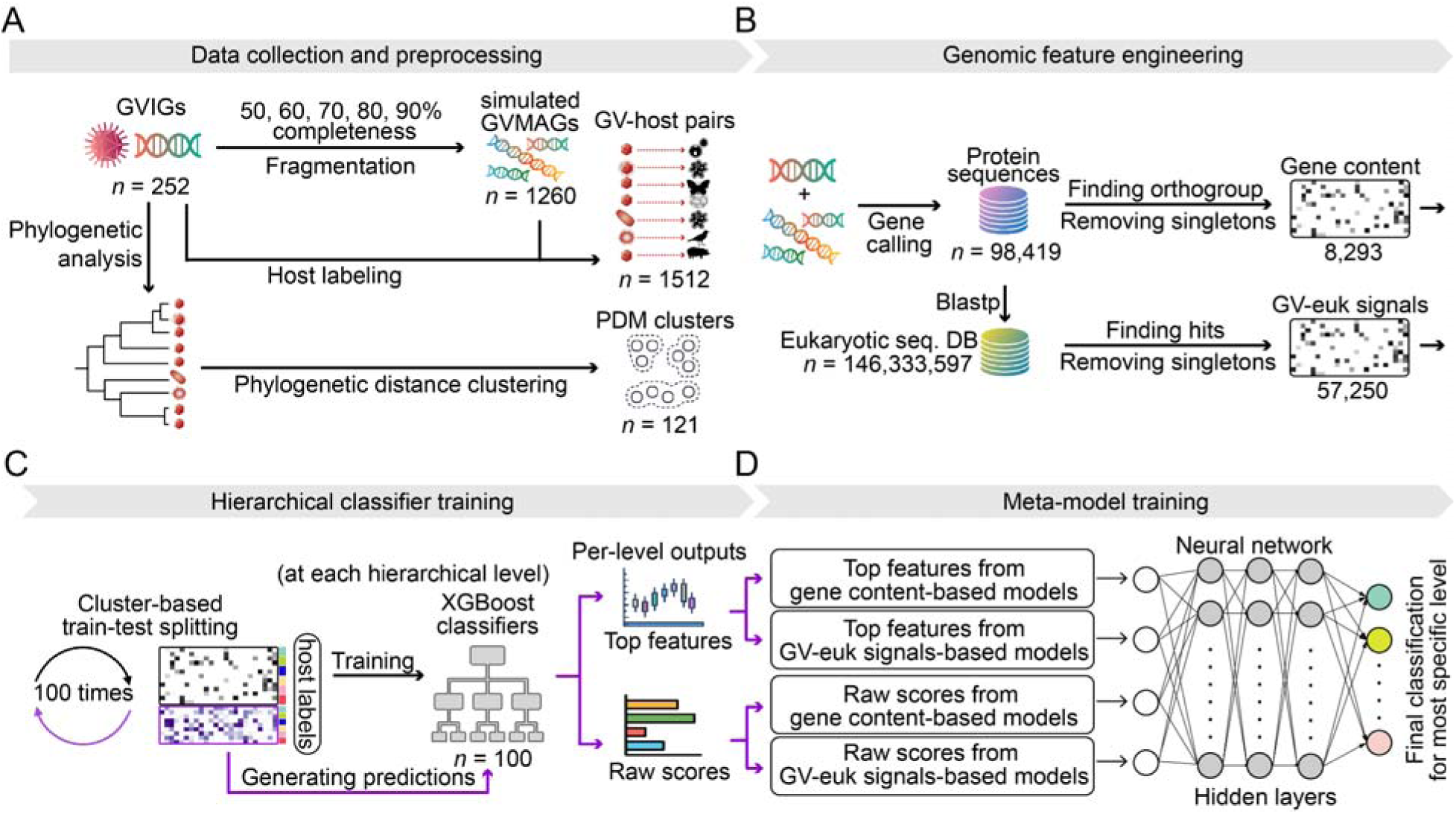
Overview of GVHoP model training pipeline. (A) Preprocessing workflow of GVIG datasets, including simulation of GVMAGs, host labeling, phylogenetic analysis, and phylogenetic clustering. (B) Feature engineering for generating feature matrices as inputs for model training. (C) Model training based on the hierarchical host labeling for the GV-host pairs. PDM cluster-based train-test splitting of the dataset was used to evaluate the model performance. Feature importance scores and raw scores were directly obtained from 100 XGBoost models. (D) Output scores from the 100 base XGBoost classifiers were used to train a neural network meta-model. Downstream host prediction at the most specific taxonomic level is determined via an *argmax* function.

**Figure 2.**
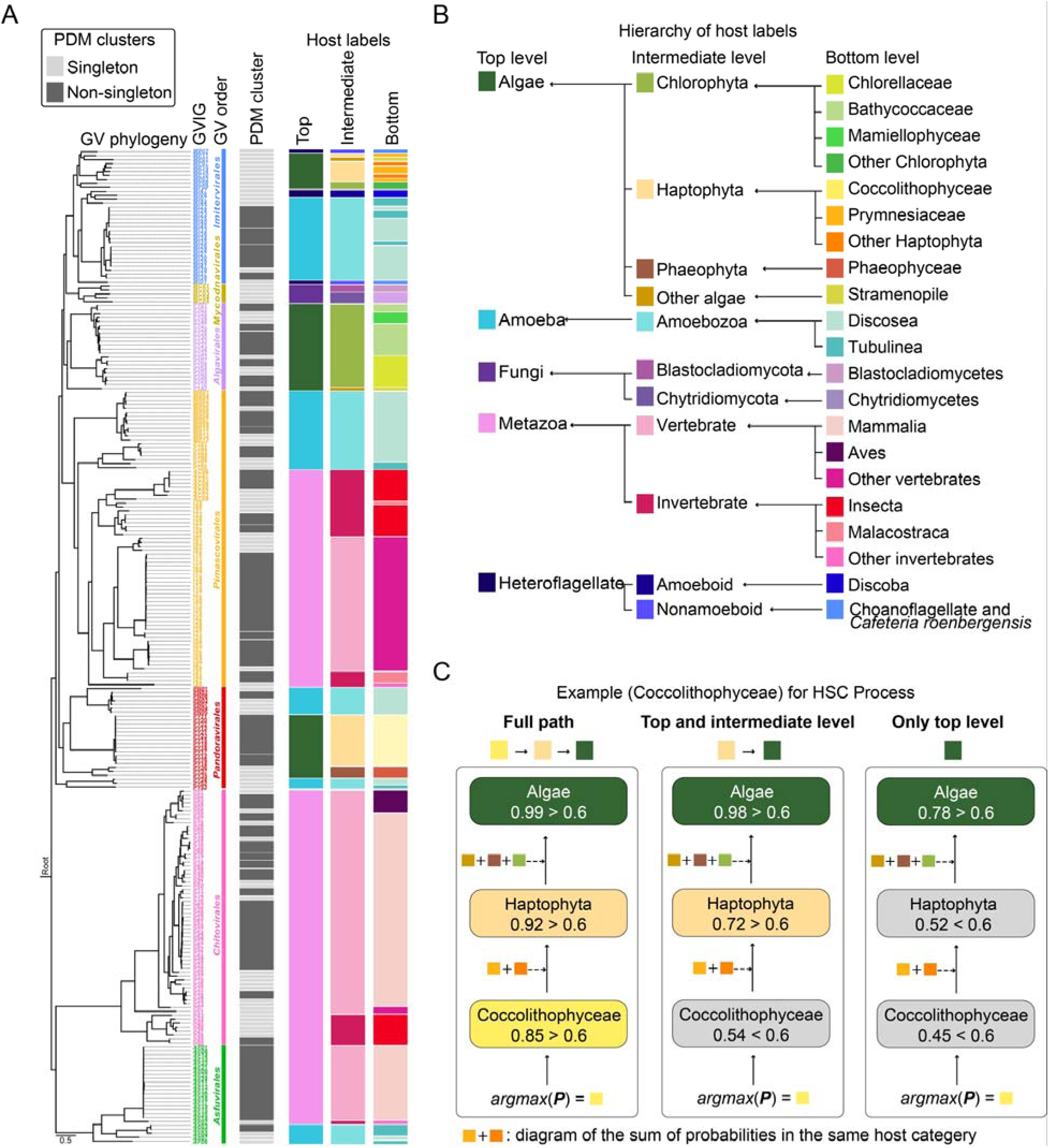
Phylogenetic clustering, host labeling, and hierarchical selective classification. (A) The phylogeny of the 252 GVIGs was inferred using a maximum-likelihood approach based on a concatenated alignment of seven GV core genes. PDM clusters with ≥ 2 GVIGs: dark gray; singleton GVIGs: light gray. (B) Ground-truth host annotations of the GVIGs at three hierarchical taxonomic levels. (C) Illustrative examples of the hierarchical selective classification (HSC) process implemented in GVHoP. Host prediction at the most specific taxonomic level is first determined using an *argmax* function. The algorithm then traverses the hierarchical path upward, aggregating the cumulative probabilities of sibling leaves sharing a common ancestor until the predefined confidence threshold is met.

Given that many GV-host relationships are represented by only one to three GV genomes, and considering the complexity of eukaryotic taxonomy, we reclassified eukaryotes into five top host types and their subordinate groups by considering both phylogenetic relationships and eco-physiological similarities (Figure 2A, 2C). This classification scheme integrates samples within underrepresented host groups and enables GVHoP to identify key features based on the physiological properties of hosts in addition to taxonomic closeness. The three-level design also allows the model to capture distinct host-associated features at more specific host levels (intermediate and bottom), thereby providing a more meaningful framework for interpreting model-identified features associated with GV-host interactions. With our objective of applying the GVHoP pipeline to all GVMAGs across ecosystems, we generated artificial genomes from each GVIG at five levels of completeness to simulate the fragmented and incomplete nature of GVMAGs (Figure 1A, Supplementary Figure S1). This dataset of simulated GVMAGs contains 1, 260 genomes, which are used for expanding the training set.

#### Genomic feature engineering and model training

We extracted three types of features from each genome to ensure comprehensive genomic representation in GVHoP. First, we clustered protein orthogroups of GVs, which are known to correlate with host diversity (54) and represent the functional potential encoded in the genome. Second, we calculated GV-euk signals to detect potential virus-host linkages through HGT. Third, we also included *k*mers, short subsequences of length *k* extracted from genomic sequences, which have been used for other virus-host predictions. For the 252 GVIGs and their 1, 260 derived, simulated GVMAGs, we generated a gene content matrix (8, 293 features), a GV-euk signal matrix (57, 250 features), and five *k*mer frequency matrices (*k* = 3-7). These matrices served as raw inputs to train base-classifiers for initial host predictions and feature extraction at different host levels (Supplementary Figure S2).

Closely related genomes can have highly similar feature patterns, which may cause data leakage when they exist in both the training and test sets. Thus, the PDM cluster-based validation scheme was applied to 121 species-level clusters, where simulated GVMAGs and their source GVIGs were assigned to the same cluster; within each iteration, these clusters were split into 70% training and 30% test sets. This splitting procedure was repeated 100 times during model training, yielding 100 trained base-classifiers for each feature matrix and host hierarchical level. While *k*mer-based classifiers had lower overall accuracy (< 80%) across 100 cluster-based 70/30 train-test splits (Supplementary Figure S2), both gene content- and GV-euk signal-based classifiers achieved high predictive performance (> 90%). Therefore, only the latter two feature sets were used for the subsequent pipeline (Figure 1B, 1C).

To integrate the predictions based on gene content and GV-euk signal features, we generated a neural-network meta-classifier that was trained on the raw prediction scores and top-ranked features from both types of classifiers (Figure 1D). Given the complex nonlinear structure observed in PCoA based on these feature sets (Supplementary Figure S3), we adopted a hybrid XGBoost-neural network approach. XGBoost first reduced the high-dimensional feature space to informative variables, mitigating overfitting, while the neural network modeled non-linear decision boundaries among the selected features. Neural-network probabilities at the most specific hierarchical level initiated the classification path, followed by a bottom-up climbing inference algorithm that traces predictions upward along the path with the highest probabilities (Figure 2D). Prediction specificity was progressively reduced until reaching a predefined confidence threshold, yielding host assignments at the highest achievable resolution supported by the model.

#### Evaluating performance of the models

The confidence threshold was optimized by systematically evaluating top-level classification performance at 0.05 probability intervals (Supplementary Figure S4A), which revealed the expected trade-off between recall and precision. To maximize detection sensitivity for diverse host groups, including underrepresented categories, while maintaining a low false discovery rate (FDR), a confidence threshold of 0.75 was selected. At this threshold and under leakage-controlled evaluation, the model achieved an overall accuracy of 0.97 while maintaining an average hierarchical coverage above 0.9 (Supplementary Figure S4B). The confusion matrices (Figure 3) show that the integrative GVHoP model achieved superior recall across most host groups compared with the single feature set classifiers. Recall is ≥ 93% for algae, amoeba, fungi, and metazoa, which are well-predicted hosts. For heteroflagellates, the most underrepresented group in the training dataset, GVHoP improved recall to 36%, compared with 12% for the gene content-based classifier, while reducing misclassification into other host groups (Figure 3). These results demonstrate that integrating gene content and GV-euk signal features provides complementary information, resulting in more robust and accurate host prediction than models relying on either feature type alone.

**Figure 3.**
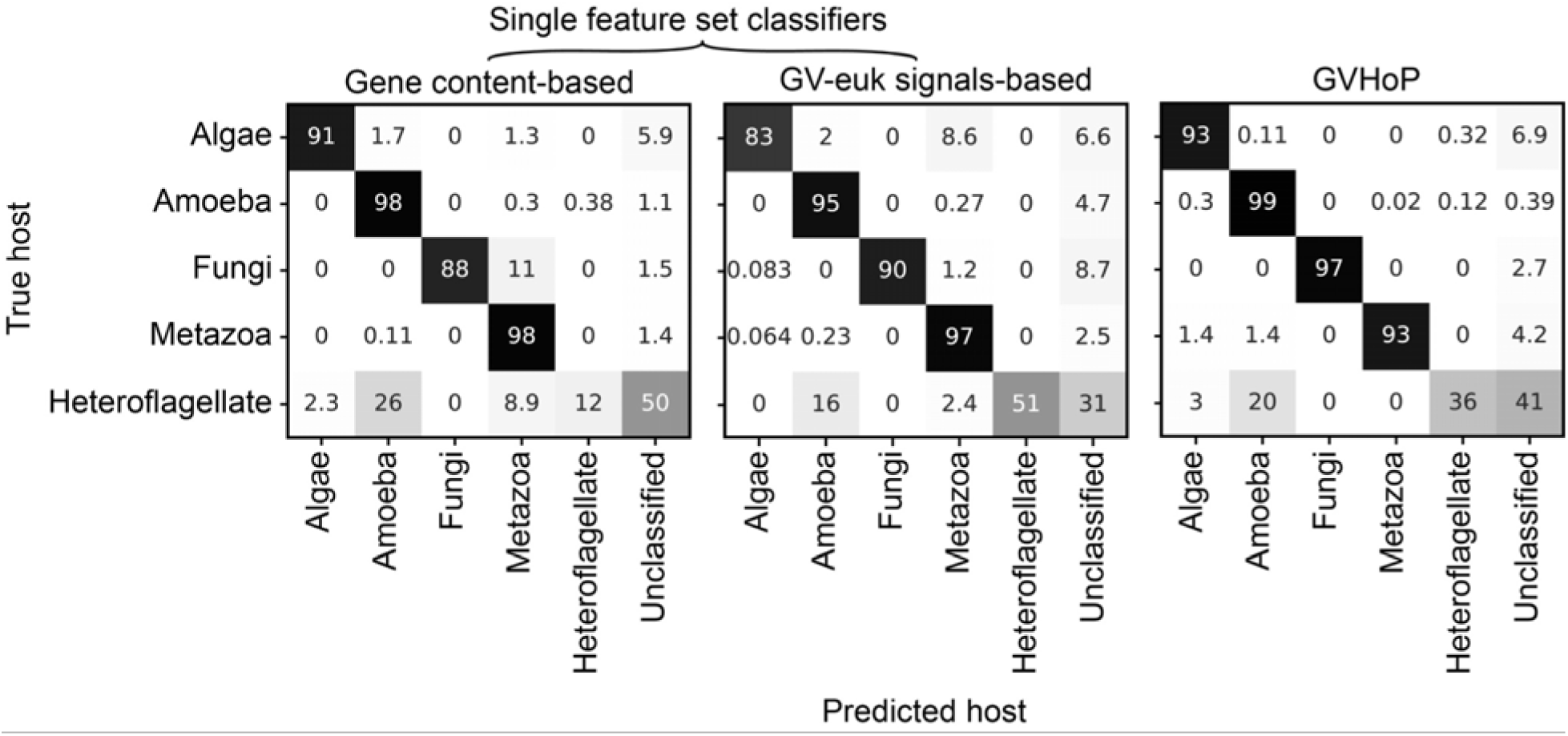
Comparison of host classification performance between single feature set classifiers and GVHoP. Classifiers based on individual feature sets (gene content or GV-euk signals) are benchmarked against GVHoP, which integrates both feature types. Each matrix displays the proportion of true hosts correctly identified within each host type, where diagonal values correspond to category-specific recall scores. The Unclassified category accounts for samples that did not satisfy the classification threshold.

### Model-informative features and their predicted functions

Feature importance scores were averaged across 100 iterations to identify informative features contributing to GV-host predictions. For each host level, the top 200 GV orthogroups (OGs) and GV-euk signals identified by the base-classifiers were extracted and deduplicated, resulting in a set of model-informative features (444 OGs and 447 GV-euk signals). To further interpret the biological context of these features, we analyzed feature prevalence across GVIGs representing five major host types and performed functional annotation of the features (Supplementary Tables 2 and 3). In total, 268 (60.4%) of the 444 top-ranked OGs and 289 (64.7%) of the 447 top-ranked GV-euk signal features (eukaryotic proteins) lack any functional annotation.

Among the 176 annotated OGs of GVIGs, predicted functions span multiple stages of the viral replication cycle (Figure 4). OGs prevalent across ≥ 2 major host types are related to genome replication, transcription, post-transcriptional regulation, protein folding, and ubiquitin-mediated regulation. DNA processing functions include replication factor C (OG00046), ATP-dependent DNA ligase (OG00061), flap endonuclease (OG00055), topoisomerase II (OG00043), DNA helicases (OG00308 and OG00586), DNA-dependent RNA polymerase subunits (OG00013 and OG00096). OGs for RNA metabolism, including mRNA capping (OG00134) and decapping enzymes (OG00112 and OG02110), RNase H (OG01000), helicase C-like proteins (OG01124), and NUDIX domain proteins (OG00152), have previously been implicated in regulating viral gene expression and modulating host antiviral responses (55–58). OGs for protein quality control and folding include DnaJ molecular chaperones (OG00082), ubiquitin carboxyl-terminal hydrolases (OG00116), and RING/RING-variant E3 ubiquitin ligases (OG00185). FK506-binding proteins (OG01751) and SWIB/MDM2 domain proteins (OG00094) have been linked to the interactions between other viruses (Influenza A virus and adenovirus) and their hosts (59, 60).

**Figure 4.**
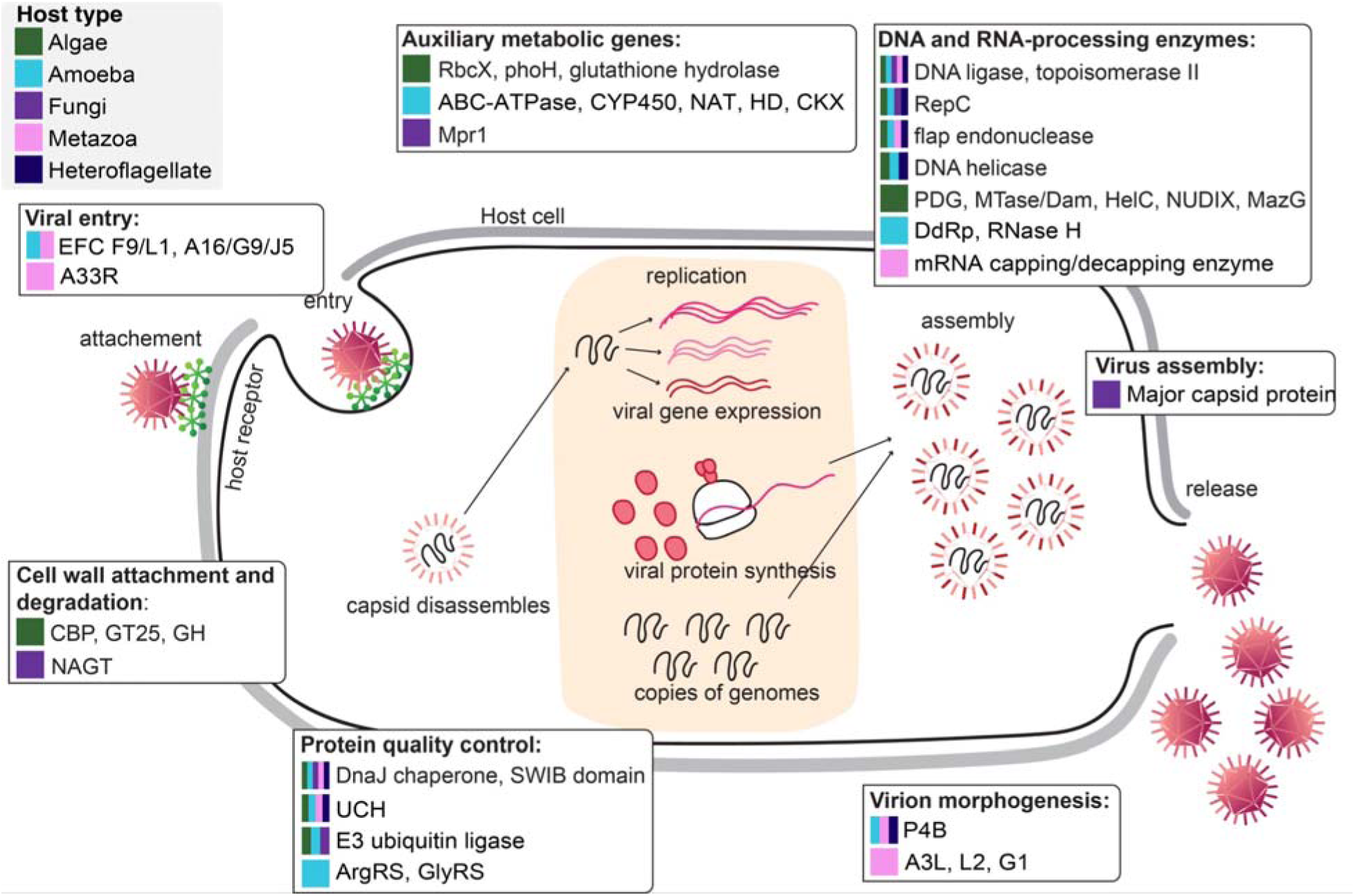
Potential functions of top model-informative features in a generalized viral replication cycle. Colors indicate major host types in which the top features were present. Description: A33R (type II integral membrane glycoprotein); ABC-ATPase (ABC transporter ATP-binding protein); ArgRS (arginyl-tRNA aminoacylation); CBP (carbohydrate-binding proteins); CKX (cytokinin dehydrogenase); CYP450 (cytochrome P450); DdRp (DNA-dependent RNA polymerase); EFC F9/L1, A16/G9/J5 (homologs of poxvirus entry-fusion complex components); GH (glycosyl hydrolases); GlyRS (glycyl-tRNA synthetase); GT25 (glycosyltransferase family 25); HD (HD-domain proteins); MazG (MazG nucleotide pyrophosphohydrolase domain); Mpr1 (histidine-containing response regulator phosphotransferase); MTase/Dam (DNA methyltransferases); NAGT (N-acetylglucosaminyltransferase); NAT (arylamine N-acetyltransferase); P4B, A3L, L2, G1 (poxvirus core proteins); PDG (pyrimidine dimer DNA glycosylase); PhoH (phosphate starvation-inducible protein); RbcX (RuBisCO chaperone); RepC (replication factor C); UCH (ubiquitin carboxyl-terminal hydrolases).

In contrast to machineries related to DNA, RNA, and proteins, top OGs annotated with other diverse functions show more distinct association with specific host types. Homologs of carbohydrate-binding proteins (OG01089), glycosyltransferase family 25 (OG01301), and glycosyl hydrolases (OG04272 and OG02738) are often associated with algal hosts (e.g., coccolithophores, *Ostreococcus*) and may modulate interactions with host surface glycans or extracellular matrices. For amoeba- and metazoa-infecting viruses, homologs of poxvirus entry-fusion complex proteins, including F9/L1 (OG00169) and A16/G9/J5 (OG00138 and OG00144), can potentially mediate membrane fusion during viral entry (61, 62). Homologs of poxvirus core proteins L2 (OG00272; hosts: mammals), P4B (OG00090; amoeba, avians, mammals, insects, and heteroflagellates [*Cafeteria*, *Naegleria*, and *Bodo*]), A3L (OG00201; avians and mammals), and G1 (OG00177; avians and mammals) have been implicated in virion morphogenesis (63–66). Homologs of autophagy-related proteins (OG00348; *Acanthamoeba* and *Naegleria*) may be involved in modulating host cell autophagy, which has been shown to be a pathway targeted by poxviruses to increase viral replication (67). Another top feature is homologs of type II integral membrane glycoprotein A33R (OG00273; mammals), which is known to be important for intracellular trafficking during poxvirus replication (68).

For algae-infecting GVs, we further identified OGs that participate in diverse metabolic processes. Examples include RbcX (OG08577; *Bathycoccus*), the assembly chaperone of RuBisCO central to photosynthesis, PhoH (OG00678; *Micromonas pusilla* and *Ostreococcus tauri*), which can regulate cellular physiology under phosphate-limited conditions, glutathione hydrolase (OG02978; *Haptolina* and *Prymnesium kappa* [Prymnesiaceae]) and alkyl sulfatase (OG07442; *Chlorella* and *Tetraselmis* green algae) for sulfur metabolism and redox balance, and pyrimidine dimer DNA glycosylase (OG00625; *Chlorella* and coccolithophores) and DNA methyltransferases (OG01607; *Chlorella* and *Ostreococcus lucimarinus*) for DNA repair and modification. Encoded by amoeba-infecting GVs, ABC transporter ATP-binding protein (OG00523), cytochrome P450 (OG00315), and arylamine N-acetyltransferase (OG00242) may modulate host nutrient transport, detoxification, redox regulation, or other pathways. HD-domain proteins (OG02752) may serve as phosphohydrolase in breakdown of nucleotides and nucleic acids. We also identified cytokinin dehydrogenase (OG01529) but its precise function in amoebae remains unclear. Additionally, arginyl-(OG00599) and glycyl-tRNA synthetase (OG02756) are among the top OGs from amoeba-infecting GVs, which often encode a larger repertoire of translation machineries (69). In fungi-infecting viruses, OG02435 has potential N-acetylglucosaminyltransferase and N-acetyltransferase functions and might be be used to regulate N-acetylglucosamine (building block of chitin) metabolism and the dynamics of fungal cell walls. OG05201, which encodes a histidine phosphotransfer protein, is potentially involved in two-component signal transduction in fungi (70).

The 447 top-ranked GV-euk signal features (eukaryotic proteins) correspond to 221 GV OGs (Supplementary Table S3), of which 55 overlap with the top GV OGs identified from the gene content feature set and are primarily involved in functions for core DNA or RNA metabolism as aforementioned. The other 131 unique and annotated OGs extracted from the top GV-euk signal features are more commonly associated with specific host types. In algae-infecting GVIGs, examples include a urea transporter (OG00906-KOO22511.1; hosts: *Haptolina*, *Phaeocystis*, and coccolithophores), which may contribute to nitrogen acquisition. A metalloprotease (OG00029-KAG5177322.1; *Prymnesium* and *Haptolina*) may participate in protein turnover. A ribonucleotide reductase (OG00024-RKP08053.1; KAL1729685.1; Coccolithophores) may support DNA synthesis by converting ribonucleotides to deoxyribonucleotides, particularly relevant to replication of large genomes for GVs.

Interestingly, we found several uncharacterized proteins encoded by GVIGs only known to infect Amoebozoa amoebae commonly used for laboratory isolation, but their top eukaryotic protein hits come from heterolobosean amoebae in another supergroup, Discoba, including *Willaertia magna* (OG00111-KAL9648785.1; *Acanthamoeba* and *Vermamoeba*), *Naegleria fowleri* (OG00111-XP_044568565.1: *Acanthamoeba*), and *Naegleria gruberi* (OG00074-XP_002674806.1: *Acanthamoeba* and *Vermamoeba*). In fungi-infecting GVIG, proteins show sequence similarity to fungal proteins involved in metabolic and biosynthetic processes, including the glycosyltransferase family (OG00300-KXS11864.1: *Allomyces*, *Blyttiomyces*, and *Synchytrium*) and the ribonucleotide-diphosphate reductase small subunit (OG00017-KAJ3035334.1: *Allomyces*, *Blyttiomyces*, *Chytriomyces*, and *Synchytrium*). An ADP/ATP carrier protein 3 identified from GV-metazoan signals (OG00024-KAJ6644940.1: Insecta, Malacostraca, Actinopteri, Amphibia, Reptilia) is potentially involved in cellular energy metabolism and mobilization of host mitochondrial deoxyribonucletide pools for viral replication (71).

### Host prediction for uncultured GVs

#### Variation in predicted hosts across environments

The largest GVMAG database to date contains 8, 508 species-level genomes across aquatic, terrestrial, and anthropogenic environments (19). To predict the hosts of the GVMAGs, we first filtered for genomes classified under the phylum *Nucleocytoviricota* and deduplicated sequences closely related to our training data using PDM clustering, which resulted in an unseen dataset of 7, 897 GVMAGs (Figure 5A). Applying GVHoP to these sequences with the threshold 0.75, we identified hosts for 5, 280, 5, 015, and 4, 425 genomes at the top, intermediate, and bottom hierarchical levels, respectively (Figure 5B). That is, GVHoP successfully predicted hosts for approximately 67% of the GVMAGs (Supplementary Table S4), thereby enhancing our biological understanding of these uncultured GVs.

**Figure 5.**
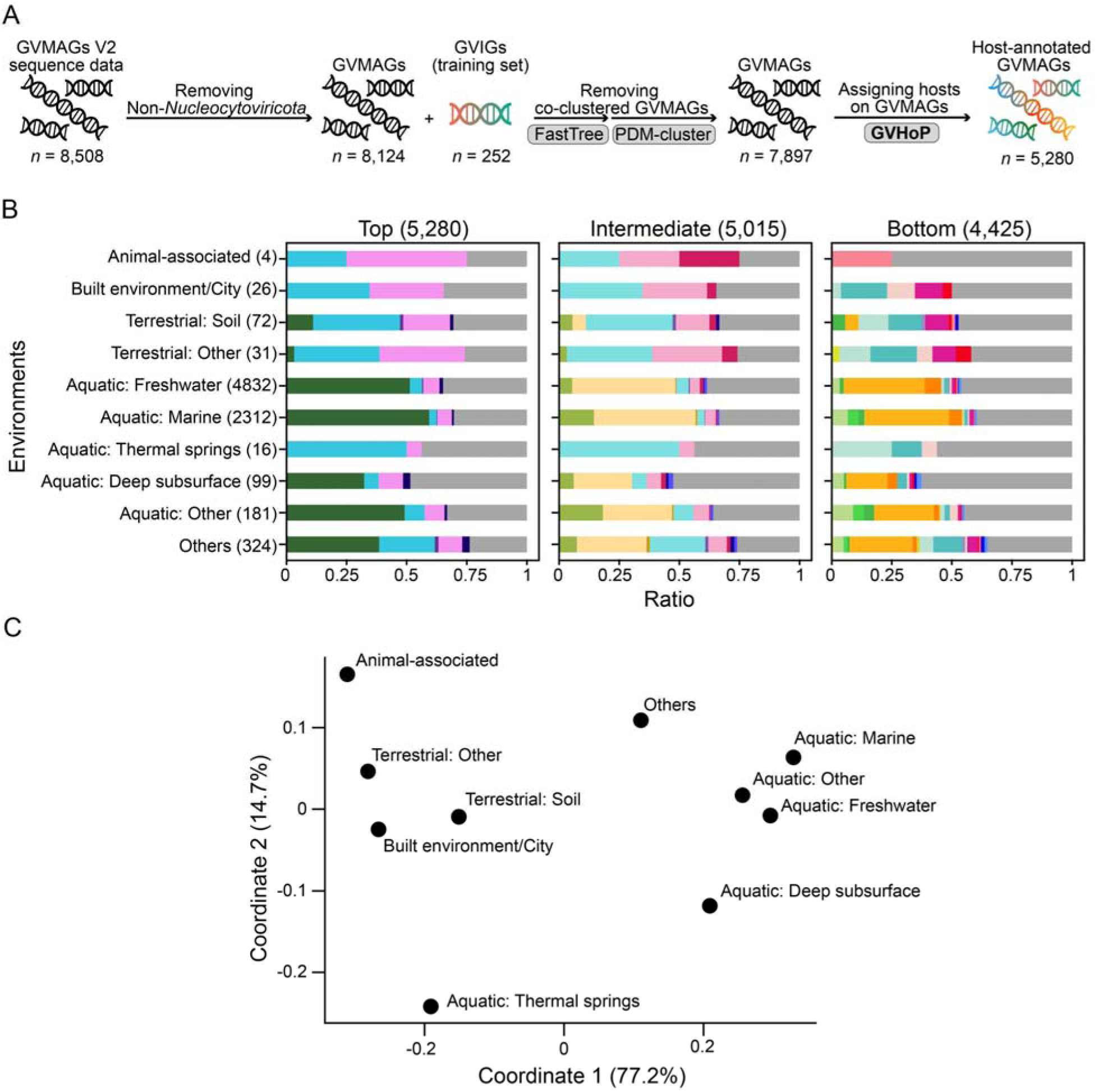
Host prediction of GVMAGs. (A) Workflow of host prediction for GVMAGs. (B) The stacked bar chart illustrates the distribution of predicted hosts of GVMAGs across different environments. The colored segments indicate the proportion of GVMAGs predicted to different host categories at three levels, which share the same color codes as in Figure 2B. (C) Dissimilarity of predicted host composition among all environments based on PCoA.

Distinct patterns of predicted host categories were observed across different environments, indicating that host composition varies with environmental context (Figure 5B). Metazoans and amoebae, respectively, represent 50% and 25% of predicted hosts in the animal-associated environment, and approximately 30% and 26% in human-built and terrestrial environments. In contrast, they account for only 5% and 4% averaged across the aquatic environments. Freshwater, marine, deep subsurface, and other aquatic environments are dominated by viruses with predicted algal hosts, which comprise 34-57% of predicted hosts, while heterotrophic flagellate hosts consistently represent a small but detectable fraction (approximately 1%). Thermal spring viromes are a notable exception among aquatic habitats, exhibiting a predicted host composition more similar to terrestrial environments, with 50% and 5% of predicted hosts being amoebae and metazoans, respectively.

Based on PCoA of pairwise dissimilarities among predicted host abundance profiles (Figure 5C; PCoA1 and PCoA2 explain 77.2% and 14.7% of the variation, respectively), we observe a major separation between aquatic and terrestrial environments. As aforementioned, thermal springs are clustered more closely with terrestrial environments than with other aquatic environments. Among the rest of the aquatic environments, we also observe a more distinct position of the deep subsurface environment.

#### GVHoP predicts hosts based on information beyond viral core-gene phylogeny

While there is broad concordance between GVHoP predictions and known hosts of each order-level clade of GVs (Supplementary Figure S5), GVHoP predicted previously unknown GV-host relationships, including *Asfuvirales* GVMAGs with heteroflagellates and algae, *Pimascovirales* with algae, *Algavirales* with heteroflagellates and metazoans, *Pandoravirales* with metazoans, *Mycodnavirales* with algae, and *Imitervirales* with metazoans (Figure 6). To evaluate the relationship between GVHoP predictions and hosts inferred purely from the viral core-gene phylogenetic relationships, we calculated prediction purity based on phylogenetic clusters derived from RED. Higher purity indicates greater agreement between GVHoP predictions and hosts known from phylogenetically close training GVIGs (Figure 7A). The mean purity increases significantly with increasing RED scores (i.e., smaller clades of more closely related GVs), with Pearson correlation coefficients ≥ 0.94 for all host levels (*p* < 0.05; Figure 7B). This is consistent with the general trend that more closely related viruses with more similar genomes tend to infect similar hosts, suggesting a purely phylogeny-based inference of hosts would only work for GVMAGs that have closely related GVIGs. However, GVHoP can predict hosts for GVMAGs irrespective of their phylogenetic relationships to known hosts. In addition, GVHoP predicted hosts of GVMAGs with low purity at high RED cutoffs (Figure 7B), which represent novel GV-host relationships that differ from the dominant hosts within their phylogenetic clusters.

**Figure 6.**
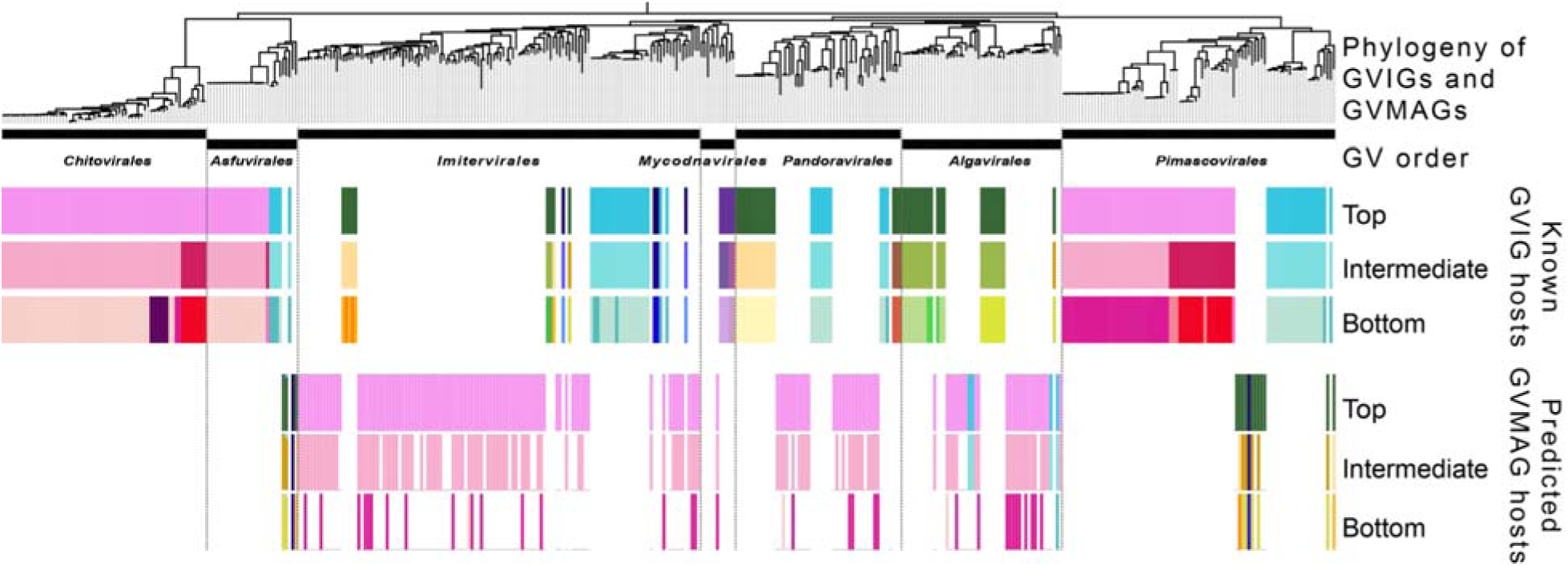
GVMAGs with GVHoP-predicted hosts distinct from known hosts of major GV orders. A phylogeny was reconstructed based on the core GV proteins (see Methods) of the 252 GVIGs and the 181 GVMAGs that were predicted to have novel hosts across GV orders. See Figure 2B for host group coloring.

**Figure 7.**
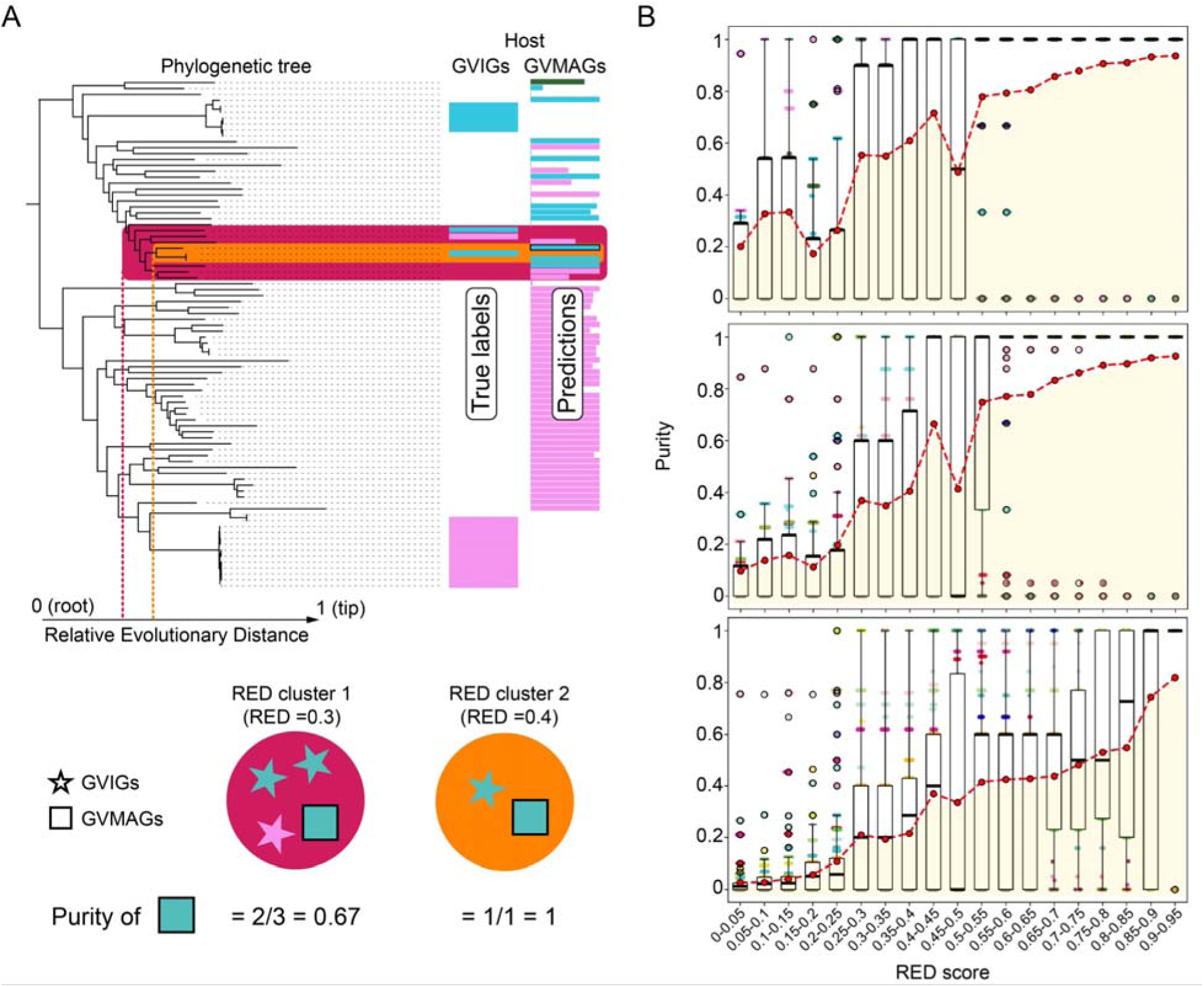
Purity of GVHoP predictions across clusters defined by different thresholds of relative evolutionary distance (RED) scores. (A) Computation of RED scores and the corresponding purity metrics for GVMAG host predictions. (B) Correlation between prediction purity and RED score at different hierarchical host levels (highest level at top). Box plots show the distribution of GVMAG purity across the RED cutoff. Each point represents a GVMAG with ≥ 1 GVIG in its RED cluster and is color-coded by its predicted host (Figure 2B). Red dots: means of GVMAG purities at each RED cutoff. Yellow shading areas highlight low-purity GVMAGs with predicted hosts distinct from their neighboring GVIGs.

## DISCUSSION

With the increasing number of GVMAGs and the high diversity of known GV hosts, the taxonomic identities of the hosts of GVMAGs have been a major knowledge gap toward understanding their roles in the environment. One way to infer potential hosts is to extrapolate the virus-host relationships from closely related GVIGs based on a phylogeny reconstructed a few core genes (35). This approach is fast and straightforward, but it is based on the assumption that the vertical inheritance and phylogenetic information of a few core genes determine the hosts. However, compared with small viruses, GV genomes are well known for having a pool of genes that are horizontally acquired and involved in virus-host interactions (5, 26, 72, 73). The phylogeny-based approach also has the limitation that it cannot be applied to newly discovered GVs that are distantly related to any lineages with known hosts. Host associations can also be inferred from co-occurrence between the marker sequences of GVs (e.g., *polB*) and eukaryotes (e.g., 18S rRNA gene) revealed through environmental sequencing (74, 75). While it can help reveal previously unknown associations between GVs and eukaryotes, this method is limited to environmental samples with sequencing datasets that have the resolution to profile the communities of both GVs and eukaryotes. It is also prone to the complex cellular, physiological, ecological, and technical factors that influence the abundances of viral and host DNA and the detection of their specific marker sequences. Here, we developed machine learning method GVHoP, which integrates viral gene content and GV-euk protein similarities, to efficiently predict the hosts of diverse, uncharacterized GVs that only have (partial) genome information. Compared with the phylogeny-based approach, GVHoP predictions are based on genome-level information rather than a few core genes that play similar roles across GVs and are less likely to determine GV-host interactions. With the design of hierarchical host labels, GVHoP enables host prediction with different resolutions that depend on the genome information available for each GV.

We reclassified the original host labels into five major host types based on phylogenetic relationships and eco-physiological similarities (Figure 2A, 2C). This broader classification provides a biologically meaningful representation of host diversity, which is unevenly represented by isolated GVs, and allows GVHoP to capture not only closely related but also ecologically similar hosts that would be overlooked in a strict taxonomic or phylogenetic scheme. With the three-level host classification, we were able to extract top features that are important for GV-host interactions across major host groups, within a major host group, or for more specific host lineages (Figure 4). These model-identified GV proteins may reflect convergent evolution associated with a broad host type (e.g., algae) or lineage-specific gene acquisitions during the adaptation to a particular eukaryotic taxon. Thus, the hierarchical classification provides a useful framework for predicting hosts of GVs and for linking genomic signatures to the biology of GV-host interactions.

We used a machine learning approach to reveal the hidden connections between the genomic information of GVs and their hosts. The PCoA based on the GV gene repertoires and GV-euk signals showed varying degrees of overlapping distributions between host types (Supplementary Figure S3). Consistent with the complex evolutionary origins and a wide variety of functions of GV genes that collectively determine GV-host relationships, GVs employ diverse strategies to infect eukaryotes that vary significantly in cell biology, physiology, ecology, and evolutionary history. For example, GVs inhabiting polar environments show diverse adaptive strategies that reflect the interplay among viral ancestry, host biology, and environmental conditions (76). It was also shown that freshwater GVs adapt to environmental changes through gene duplications, HGTs, and positive selection, particularly in genes associated with host interactions (75). Similarly, highly dynamic genomic regions enriched in host-interaction genes can promote genome diversification and host adaptation through rearrangement and genetic exchange with both eukaryotic and bacterial genomes (77). Together, these patterns highlight the combined influences of viral evolution, genome plasticity, host interactions, and environmental adaptation on GV-host relationships and the need for a machine learning framework for GV host prediction.

A core component of GVHoP is the hierarchical bottom-up climbing inference strategy (Figure 2D), which, to our knowledge, represents the first application of hierarchical inference to host prediction. Rather than making predictions at a fixed taxonomic resolution, this approach with our reclassified host types allows GVHoP to adjust the resolution of host assignment according to the strength of the available information. Compared with phage-host predictors that can benefit from abundant host information derived from genome annotations and publications (20–22, 24, 31, 78), GV host prediction across divergent eukaryotes faces the challenge of lacking reliable host labels, limited training data, and increasing uncertainty at finer taxonomic resolution. In GVHoP, uncertainty is explicitly incorporated into the prediction process. Predictions with sufficient information can be resolved to more specific host groups, whereas those with weaker support are retained at broader taxonomic levels, reducing the risk of overinterpreting uncertain host assignments. The improved recall observed for challenging groups such as heteroflagellates further indicates that hierarchical inference can recover biologically meaningful signals that may be difficult to capture using conventional flat classification (Figure 3).

GV gene content and GV-euk signals capture different aspects of GV genome evolution. Gene content provides information about the repertoire of viral genes and functions, whereas GV-eukaryote signals capture potential evolutionary connections between viral and host genomes through past HGTs. By combining these two types of features that complement each other, GVHoP provides a more comprehensive representation of host-associated information. This is evident from the improved performance of the integrated model (Figure 3), as well as the distinct genes that were identified as top features of each set for determining GV-host relationships. In terms of biological functions, we see that genes involved in core information processing central to viral replication can be potentially important genes for GV-host relationships across distantly related eukaryotic hosts, whereas genes with more specific metabolic or physiological functions tend to be associated with specific host lineages. It should be noted that many highly informative genes lack known functions, suggesting they can be top candidates for further functional investigation into their roles in GV biology. While GVHoP predictions of GVMAG hosts show a certain level of agreement with the known hosts of closely related GVIGs, in particular when they are highly similar (Figure 7), GVHoP also predicted hosts of GVMAGs that are distinct from known hosts, which lead to low-purity cases (Figure 6, 7B) and may represent previously unrecognized hosts of established GV orders. With more GVs isolated and their hosts identified in the future, these predictions can be further tested. In particular, we see predictions of algae, metazoans, and heteroflagellates as novel hosts for GV orders that so far only have amoebae or other hosts, which implies there are more host shifts during GV evolution than recognized so far.

Viruses are ubiquitous across diverse ecosystems and regulate microbial communities, nutrient and carbon cycling, genetic exchange, and host metabolism through auxiliary metabolic genes and HGTs (79). The application of GVHoP to environmental GVMAGs revealed clear differences in predicted host composition among different environments. Human-built and other terrestrial environments are enriched in GVs with predicted metazoan and amoebal hosts, whereas aquatic environments show a greater contribution of predicted algal hosts (Figure 5B). Previous study has shown that factors such as salinity and the aquatic-terrestrial divide constrain viral distributions and select for viral adaptations that correspond to the ecological adaptations of their hosts, thereby limiting viruses to specific ecological niches (80). Given the close ecological and evolutionary coupling between GVs and their hosts, variation in host communities likely has direct influence on virome structure and contributes to the environment-specific patterns observed in viral assemblages. The separation of environments based on predicted host composition therefore provides a potential link between environmental context, host community structure, and GV diversity (Figure 5C). Overall, these model-predicted patterns are consistent with a role for environmental filtering of host communities in structuring GV-host associations. However, it merits further investigation into the role of other factors such as the productivity and environmental stability of virions, which can also potentially affect the composition of GVs in a specific environment.

In summary, our results demonstrate that GV-host prediction can benefit from integrating multiple genomic signals and explicitly accounting for the hierarchical structure and uncertainty of host classification, leading to improved prediction for diverse, unevenly represented eukaryotic host groups. GVHoP enabled host prediction for a large fraction of uncultivated GVs, assigning candidate hosts to 5, 280 GVMAGs. The machine learning pipeline also identified both functionally annotated and uncharacterized genes that can play key roles in determining GV-host interactions. By integrating gene content, GV-eukaryote signals, and hierarchical inference, GVHoP extends host prediction beyond conventional marker- and phylogeny-based approaches and provides a framework for exploring previously unresolved GV-host relationships at a large scale.

## ACKNOWLEDGEMENTS

We are grateful to Hsin-Chou Yang, An-Chi Wei, Sen-Lin Tang, and Yu-Wei Wu for their support and discussions. The work conducted by the U.S. Department of Energy Joint Genome Institute (https://ror.org/04xm1d337), a DOE Office of Science User Facility, is supported by the Office of Science of the U.S. Department of Energy operated under Contract No. DE-AC02-05CH11231.

## AUTHOR CONTRIBUTIONS

Hsin-Ying Chang: Conceptualization, Data Curation, Formal analysis, Software, Investigation, Methodology, Visualization, Writing – original draft. Frederik Schulz: Conceptualization, Investigation, Writing – review & editing, Funding acquisition, Supervision. Chuan Ku: Conceptualization, Investigation, Writing – review & editing, Funding acquisition, Resources, Supervision, Project administration.

## COMPETING INTERESTS

All authors declare no competing interests.

## FUNDING

This research was funded by the Academia Sinica grant AS-CDA-110-L01 and the National Science and Technology Council, Taiwan grants NSTC 114 2628 B 001 012 and 115-2628-B-001-007 (C.K.).

## DATA AVAILABILITY

The custom scripts, GVOG sequences, databases, and GV-host predictive models are available on Zenodo at https://doi.org/10.5281/zenodo.22199562. The GVHoP software is available at https://github.com/hyhazelchang/GVHoP.

## Notes

### Competing Interest Statement

The authors have declared no competing interest.

