## Supplementary material for "Machine learning prediction of eukaryotic hosts for giant viruses": Figure S1-S5

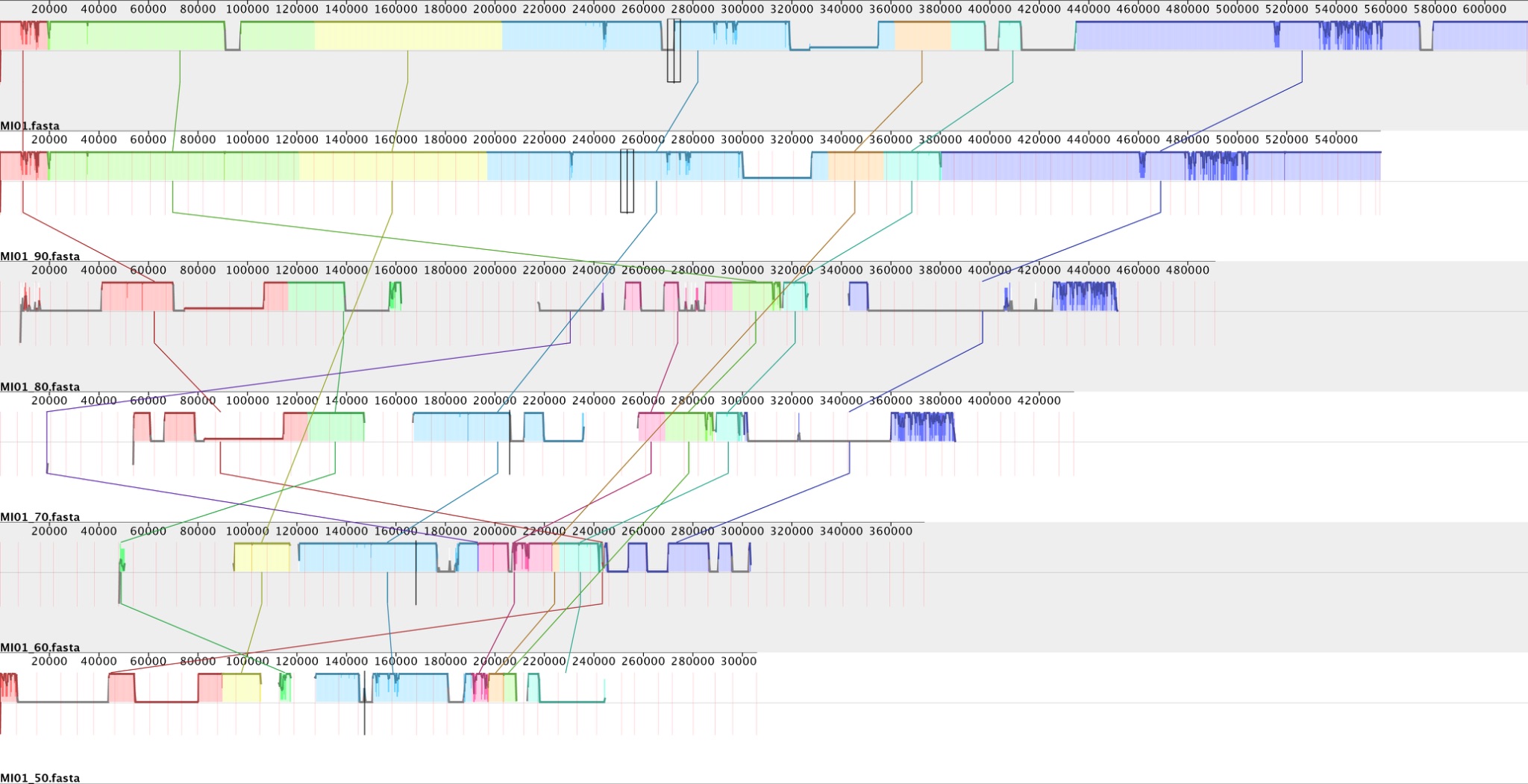

**Figure S1.** Sequence alignment of artificial giant virus metagenome-assembled genomes featuring different completeness levels (90%, 80%, 70%, 60%, and 50%).

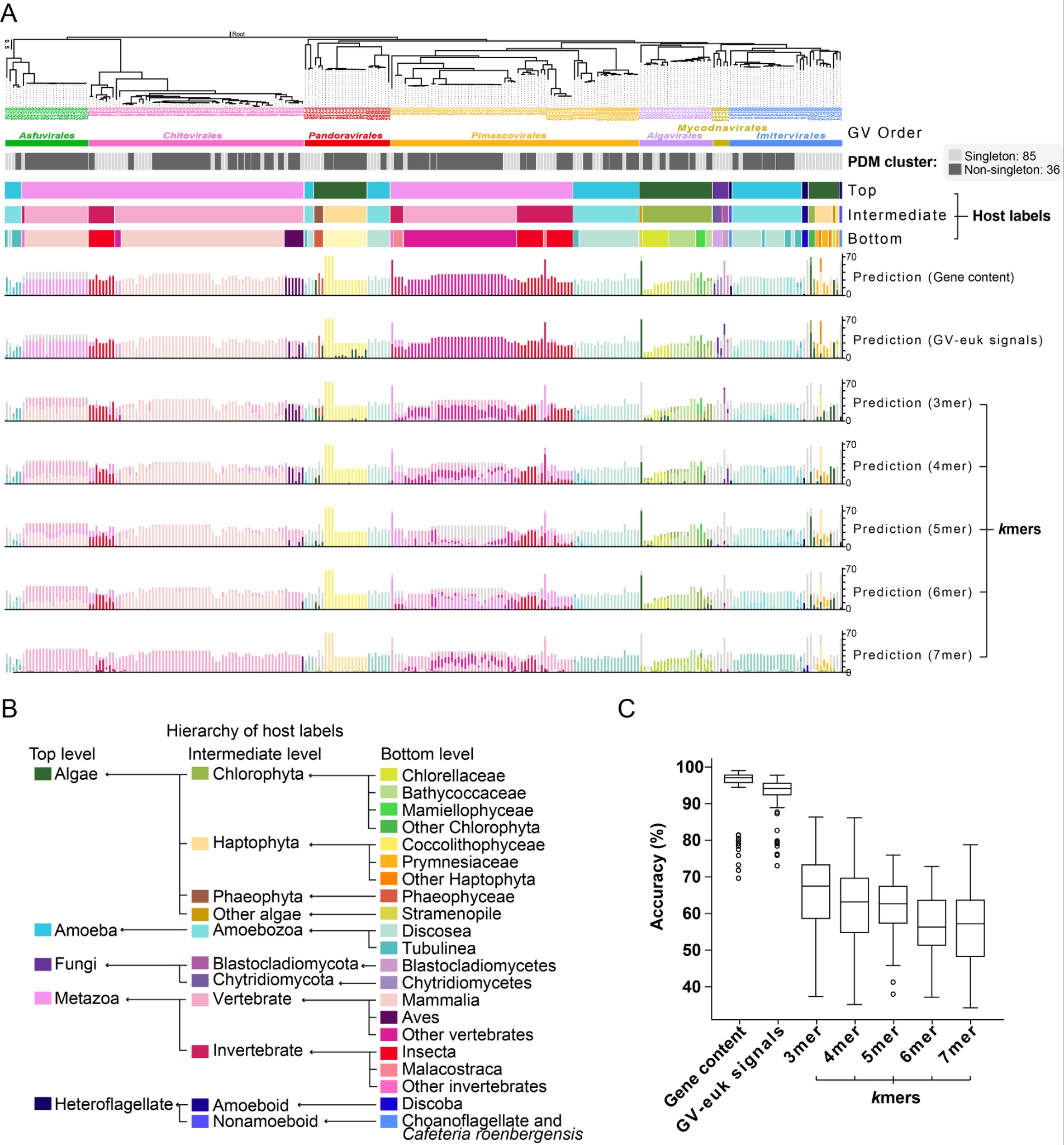

**Figure S2. Comparison of host prediction performance across different genomic feature sets.** (A) Alignment of ground truth host labels and model predictions with viral phylogeny. The top panel shows the phylogenetic tree of GVs annotated by GV order and PDM cluster. Below the tree, color-coded tracks indicate the ground truth host labels across three hierarchical levels (Top, Intermediate, Bottom). The bottom bar plots show sample-level prediction across 100 models trained on gene content, GV-euk signals, and *k*mer frequencies (*k*=3-7). (B) Hierarchical structure of host taxonomic labels used for prediction, showing the relationships among Top, Intermediate, and Bottom levels. (C) Overall prediction accuracy (%) compared across feature sets. Box plots show the median (center line), and outlier data points.

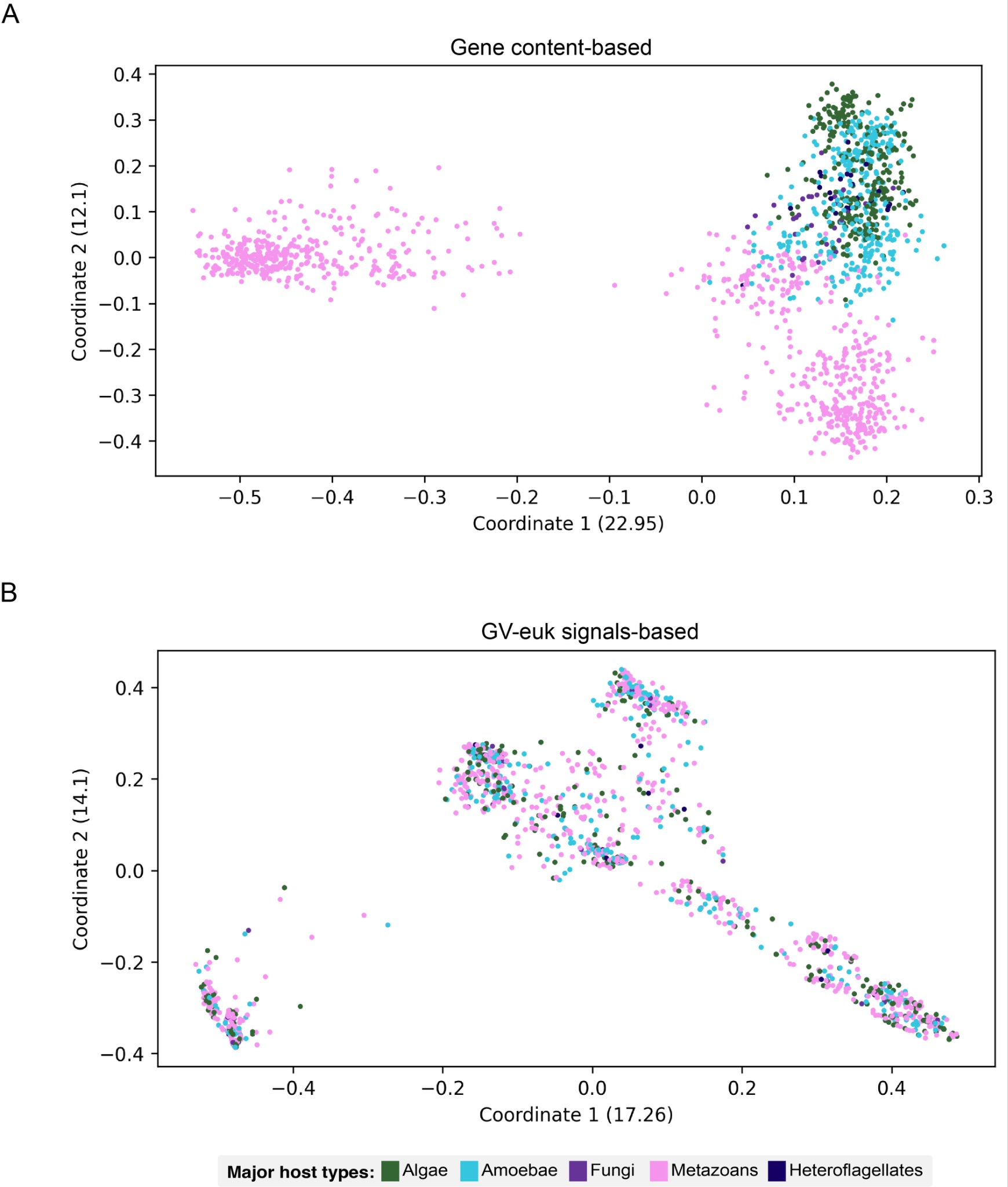

**Figure S3. Principal Coordinate Analysis (PCoA) of genomic dissimilarity based on GV gene content and GV-euk signals.** Each dot indicates a GVIG or a simulated GVMAG. (A) Gene content-based: Ordination plot based on gene content dissimilarity. (B) GV-euk signals: Ordination plot based on GV-euk signals.

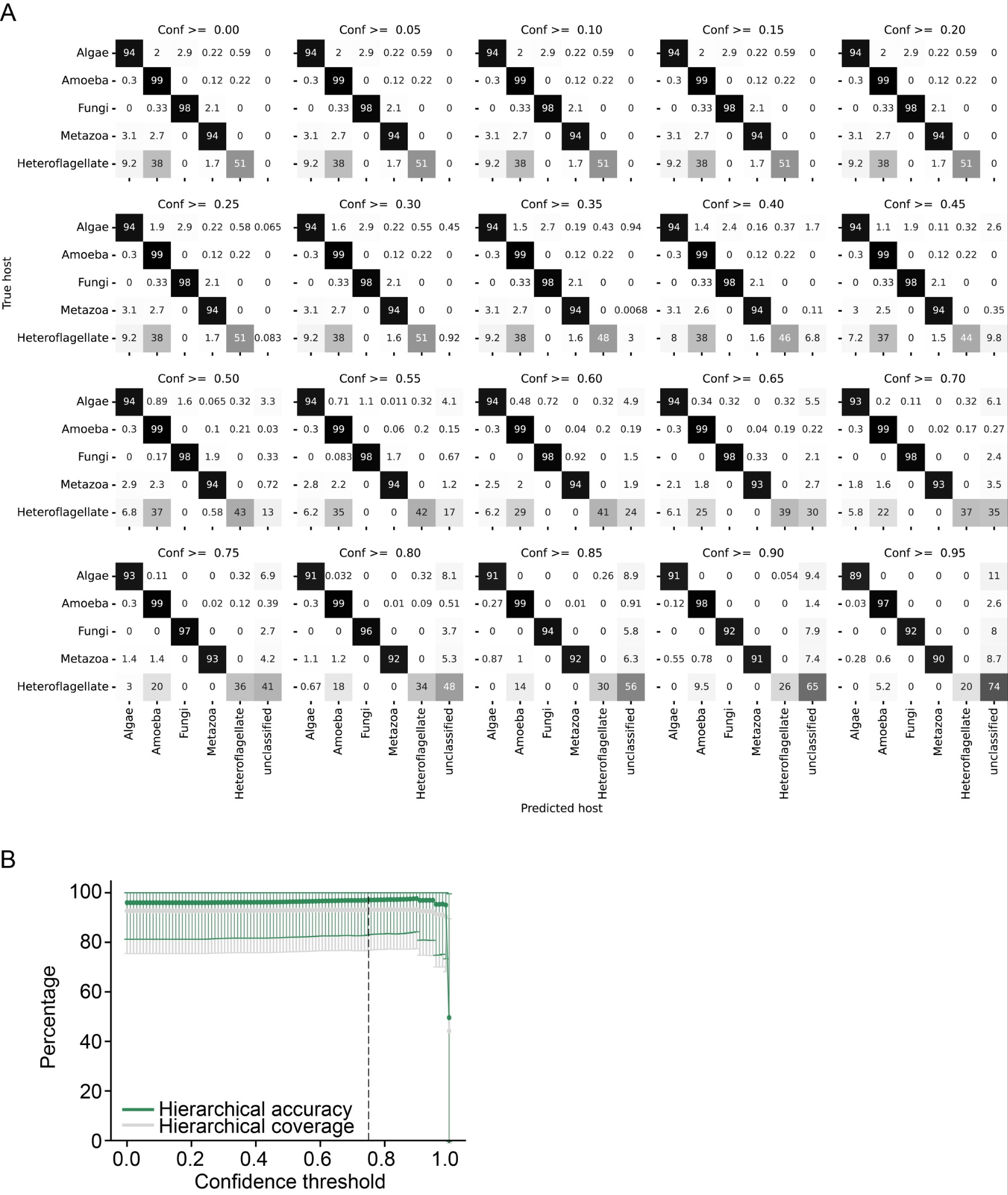

**Figure S4. Performance metrics of GVHoP.** (A) Confusion matrices for the top-level classes are presented at progressively stricter confidence thresholds (0.05 increments). The displayed data correspond to calibration performed across all PDM clusters. The values within each cell show the proportion of predicted hosts, as well as threshold-based unclassified outputs, relative to the corresponding true host. (B) Hierarchical accuracy (green) and coverage (grey) vs. confidence threshold. The dashed vertical line marks the 0.75 operating point. Error bars represent the standard deviation across leave-one-cluster-out cross-validation folds.

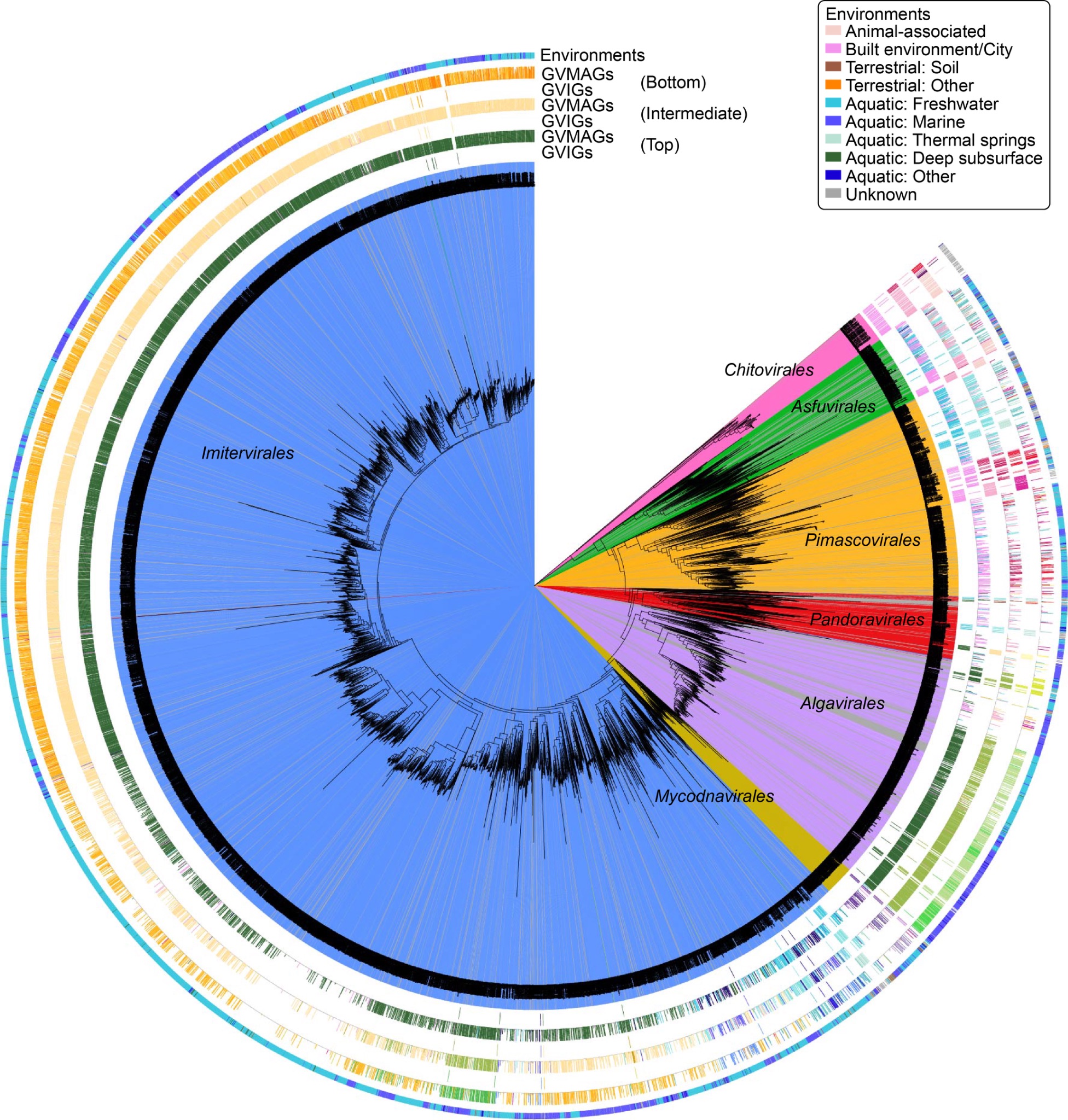

**Figure S5. Overview of known hosts of 252 GVIGs and predicted hosts of 7,897 GVMAGs, and environments of GVMAGs.** Background shading indicates the different GV orders on the phylogenetic tree. The surrounding concentric tracks are organized from inside to outside as follows: tracks 1, 3, and 5: host taxonomic hierarchy of isolated genomes, ordered from highest to lowest taxonomic rank; tracks 2, 4, and 6: predicted host taxonomy for GVMAGs at each corresponding level; outermost Track: source environments from which the GVMAGs were recovered. See Figure S2B for host group coloring. Color schemes for environments of GVMAGs are shown in the upper-right corner.
